# FastIntegration: a versatile R package for accessing and integrating large-scale single-cell RNA-seq data

**DOI:** 10.1101/2022.05.10.491296

**Authors:** Mengwei Li, Xiaomeng Zhang, Kok Siong Ang, Jinmiao Chen

## Abstract

Constructing atlas-scale comprehensive cell maps from publicly available data enables extensive data mining to achieve novel biological insights. Data integration to construct such maps require both harmonized datasets and an atlas-scale capable data integration tool. The first requirement is met by DISCO, a comprehensive repository of harmonized publicly available single cell data with standardized annotation. To meet the second requirement, the tool must have the capacity to integrate hundreds if not thousands of samples within an acceptable time frame. Moreover, it should output batch-corrected gene expression values to facilitate downstream analyses. Here, we present FastIntegration, a package which allows users to access and integrate public data in a convenient way. FastIntegration provides a fast and high-capacity version of Seurat Integration which can integrate more than 4 million cells within 2 days. It outputs batch corrected values for all genes that we can use for downstream analyses. For the first time, we demonstrated that using batch corrected values can improve the performance of downstream analyses. In particular, we found more accurately identified differentially expressed genes for cell types that are not shared between batches. Moreover, we also showed that FastIntegration outperforms existing methods for both homogeneous and heterogeneous data integration. FastIntegration also provides an API for programmatic access to data hosted on DISCO, a single-cell RNA-seq database that contains more than 5200 single-cell datasets. Users can filter for data at the sample level by tissue, disease and platform etc., and at cell level by specifying the expressed and unexpressed genes.

## Background

Single-cell omics technologies are advancing at a rapid pace, with the data throughput still growing from new techniques being developed^1,2^. Using these tools, researchers are generating ever larger datasets and uncovering new cell subpopulations in both healthy and diseased tissues. By integrating data across studies, the power and resolution of single cell studies are greatly increased to enable new discoveries^3,4^ Integration of publicly available data into atlases is particularly needed to construct consensus reference maps, together with batch corrected gene expression values that can be used for various downstream analyses. However, existing batch integration methods suffer from different drawbacks, namely being unable to scale up to a large number (hundreds or even thousands) of samples and millions of cells, or unable to produce batch corrected values^5–9^.

In our published benchmarking study^10^, wng new cell subpopulations in both healthy and diseased tissues. By integrating data across studies, the power and resolution of single cell studies are greatly increased to enable new discoveries^3,4^. Integration of publicly available data into atlases is particularly needed to construct consensus reference maps, toge found Harmony, LIGER, and Seurat v3 to be the top performing methods for batch integration, while Luecken et al. found scANVI, Scanorama, scVI, and scGen^11^ to be superior for atlas-level data integration. Except for Seurat and Scanorama, the other methods do not return batch corrected values. Furthermore, scGen and scANVI require cell type information to guide the integration, which is often not available. Though Seurat Integration returns batch corrected values, it cannot handle large-scale data integration with about 100K cells being the upper limit. Moreover, Seurat by default integrates only 2000 highly variable genes (HVGs) and returns their corrected values. If all available genes are used, it encounters performance issues when integrating more than 30 samples. Meanwhile, it remains hotly debated whether batch corrected values or uncorrected values should be used for postintegration downstream analyses such as identification of differentially expressed genes (DEGs), inference of regulon activities, and cell-cell interactions. Here, we present FastIntegration that performs efficient integration of hundreds and even thousands of samples, and outputs batch-corrected values for all genes. Furthermore, we demonstrate that the batch-corrected values improve performance in downstream analyses including DEG identification and transcription factor activity quantification.

## Results and Discussion

### FastIntegration significantly improves time efficiency and memory usage over Seurat V4 and enables large-scale data integration

We created FastIntegration by improving the original Seurat batch integration functions in several aspects (Figure 1A, details given in Methods). Most importantly, FastIntegration performs principal component analysis (PCA) only once and employs the resulting latent space during the iterative batch correction. By avoiding PCA re-computation at each iteration, the gene expression correction step can be parallelized and sped up. We added outlier detection and remove them at each round of integration to prevent overcorrection. We also optimized the use of variables to reduce memory usage and parallelized various functions to improve overall runtime performance.

**Figure 1:**
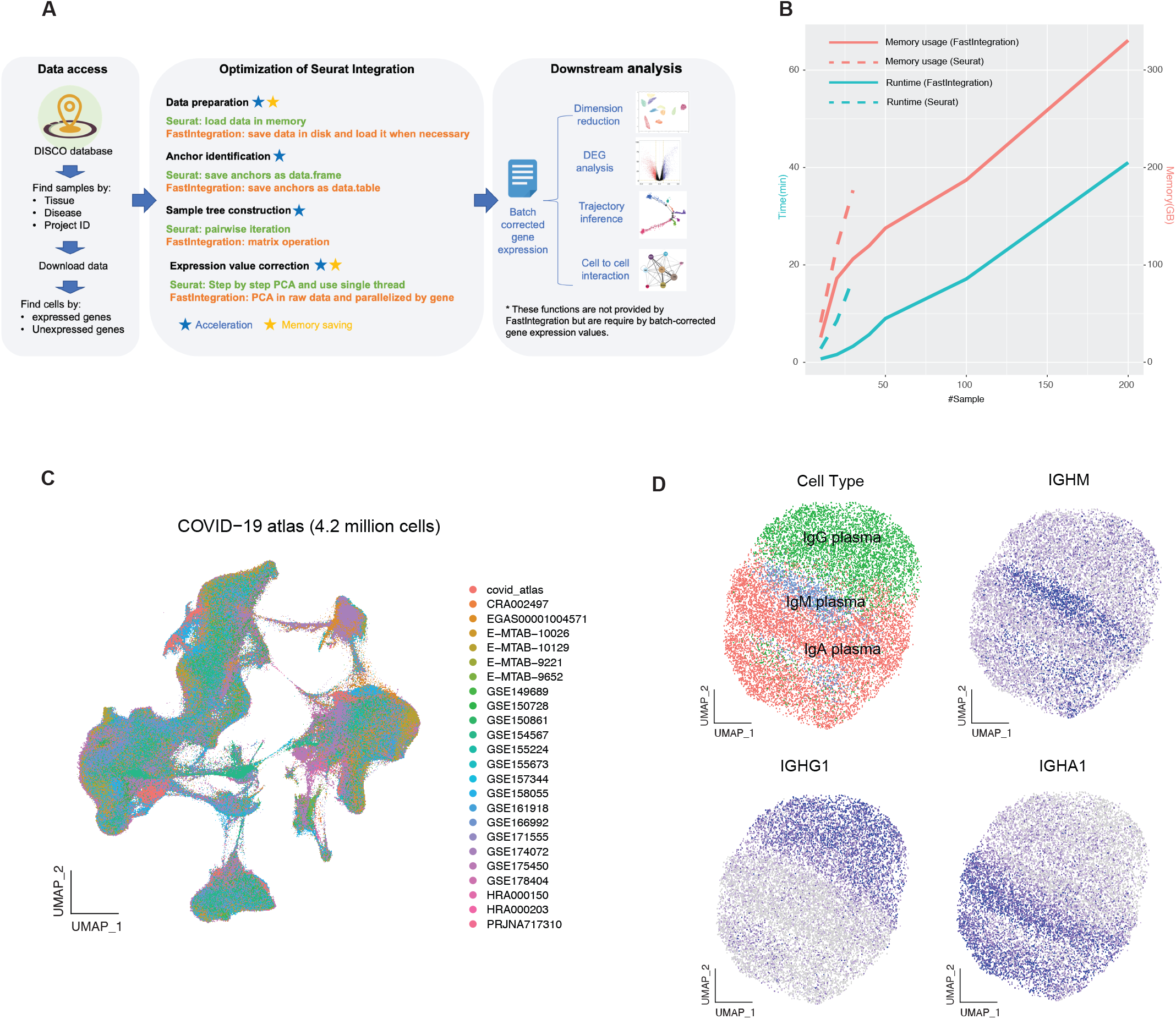
Comparison between FastIntegration and Seurat. A) The framework of FastIntegration. B) Computational efficiency benchmarks of Seurat and FastIntegration. C) UMAP of COVID-19 atlas, encompassing 4.2 million cells from 877 blood samples. D) UMAP and feature plots of data integrated plasma cells.

As we intend FastIntegration to be a fast and high-capacity version of Seurat integration, we first assessed the results produced by these two methods for consistency with the same input dataset of 20 blood samples. Both methods obtained high Adjusted Rand Index (ARI) scores based on the ground-truth cell type labels and were highly consistent between themselves with clusters showing high mutual ARI scores (Figure S1). In this dataset, neutrophils only appear in two batches, posing a challenge to batch integration methods to maintain them separate from other similar myeloid cells. Seurat mixed the neutrophils with monocytes while FastIntegration was able to retain them as a distinct cluster, which can be attributed to our overcorrection prevention steps (Figure S1A-B). Next, we compared their memory and time usage for integrating 10 to 200 samples (Figure 1B). When integrating the same number of samples, FastIntegration required less memory and time than Seurat. Moreover, Seurat was unable to integrate more than 30 samples, showing a “Cholmod error” caused by a failure to create a large sparse matrix. In contrast, FastIntegration was able to integrate 200 samples within 40 minutes with 100 threads.

To further demonstrate the large-scale integration capabilities of FastIntegration, we integrated 4.2 million cells from 877 blood samples collected from COVID-19 patients and healthy donors. The integration took less than two days with 150 threads and 2 TB peak memory usage. After integration, inter-study variations were removed (Figure 1C) while the major cell types were well separated, confirmed by their marker gene expressions (Figure S2). Strikingly, when we extracted the plasma cells and re-clustered them, we could clearly distinguish between IgG, IgM, and IgA expressing plasma cells (Figure 1D), which demonstrated that the integrated data retains the subtle differences between cells. We also applied FastIntegration on the DISCO database (https://www.immunesinglecell.org/)^12^ to create integrated atlases for different tissues, diseases, and cell types.

### FastIntegration performs well on both homogeneous and heterogeneous data integration

Current benchmarking studies^10,11^ have focused on the performance of integrating homogeneous datasets derived from the same tissue type. With the exponential growth of single-cell data, integration of heterogeneous datasets across different tissue types is becoming increasingly common. For example, several studies have integrated blood with lung tissue samples from COVID-19 patients to study the systematic immune responses at different sites^13,14^. Here, we evaluated the performance of FastIntegration and four other state-of-the-art methods, namely Harmony^6^, BBKNN^7^, scVI^8^, and Scanorama^9^ in integrating large numbers of homogeneous and heterogeneous datasets. The homogeneous datasets comprised of 50 blood samples, and the heterogeneous datasets consisted of 50 blood and 10 lung samples. Sample Local Inverse Simpson’s Index (sLISI) and inverse cell type LISI (1/cLISI) scores were used to assess batch mixing and cell type separation respectively, with a higher value denoting better performance. As BBKNN outputs only a graph, we could not calculate its LISI scores. Adjusted rand index (ARI), a global evaluation metric, was used to assess the concordance between clustering and manual cell type annotation.

For both homogeneous and heterogeneous data integration, FastIntegration and BBKNN produced the highest ARI scores (Figure 2A). However, BBKNN returns a distance-based neighbor list only, which prevents some downstream analyses. In terms of sLISI scores, Harmony was the top method with FastIntegration ranked second for both homogenous and heterogeneous integration (Figure 2B, Figure S3). In terms of cLISI, FastIntegration was the best method for both homogenous and heterogeneous integration (Figure 2B, Figure S3). Among the evaluated methods, FastIntegration struck the best balance between cell type separation (cLISI) and batch mixing (sLISI) (Figure 2B). In the homogeneous dataset, only 2 of 50 samples contained neutrophils, and only scVI, Scanorama, and FastIntegration were able to maintain the neutrophils as a separate cluster (Figure S4). This means that they are less likely to overcorrect and can better maintain separation of batch-specific cell types than Harmony and BBKNN. For the heterogeneous dataset, only FastIntegration was able to integrate the immune cells from both blood and lung samples (Figure S5). Collectively, FastIntegration showed superior and stable performance for both homogeneous and heterogeneous data integration.

**Figure 2:**
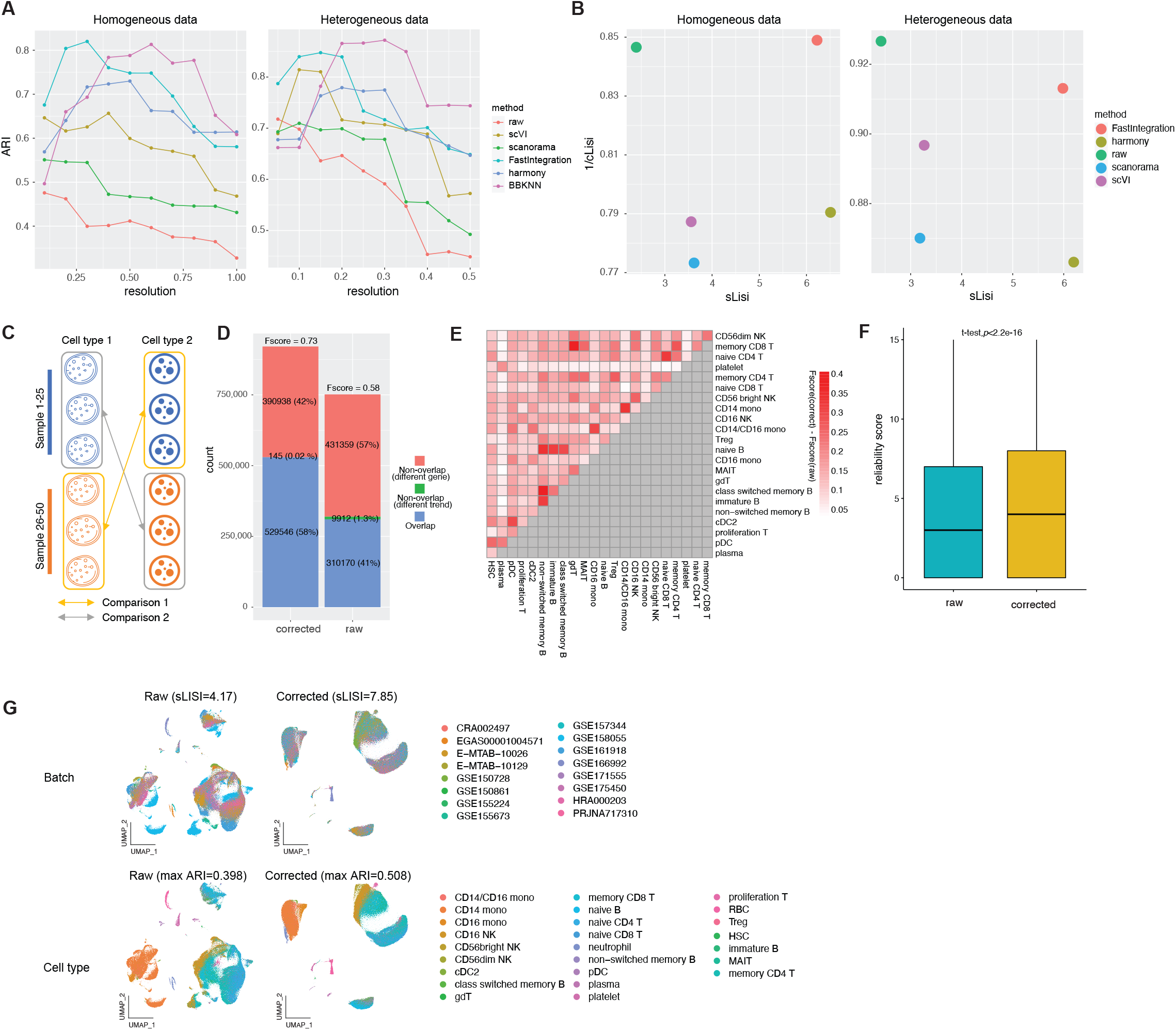
Qualitative evaluation of FastIntegration and batch-corrected gene expression value. A) ARI and B) LISI of different data integration methods for integrating the homogeneous (left) and heterogeneous (right) datasets. C) The strategy for DEGs identification. D) Conservation of DEGs identified when using raw and corrected values. E) The differences in F-scores between raw and corrected values for each cell type. F) The frequency of each DEG found in individual samples (conservative score). G) UMAPs generated with TF activities computed from the raw (left) and corrected (right) values.

### Batch corrected gene expression values generated by FastIntegration can be used for downstream analyses

The batch correction functions of FastIntegration and Seurat aim to align shared cell types across batches, removing batch effects present in the gene expression values in this process. The resulting batch-corrected values are potentially usable for various downstream analyses but there has been no study that systematically assessed the performance of batch corrected values in downstream analyses. Moreover, Seurat integration by default uses the top 2000 highly variable genes and returns the corrected values only for these genes. If all available genes are used, it encounters performance issues when integrating more than 30,000 cells. As a result, using “FindConservedMarkers” on the uncorrected values is recommended for postintegration Differential Gene Expression (DGE) analysis (https://github.com/satijalab/seurat/issues/1717). However, FindConservedMarkers cannot identify DEGs of clusters that only exist in a subset of batches. Given that FastIntegration can integrate thousands of batches and output batch corrected expression values for all genes, we evaluated the differences between using batch-corrected and raw values in DEG identification and quantification of transcription factor (TF) activity.

The homogenous dataset of 50 blood samples was used to assess DEG identification. For each cell type pair, we identified their DEGs twice using different sample subsets. The first batch of DEGs was identified between cell type 1 in samples 1 to 25, and cell type 2 in samples 26 to 50, and the second batch was identified between cell type 1 in samples 26 to 50, and cell type 2 in samples 1 to 25 (Figure 2C). This mimics the situation where some cell types only exist in a subset of samples. We found that the percentage of overlapping DEGs identified using the corrected values is much larger (58% vs 41%) than those identified using the uncorrected values, implying that the DEGs identified using the corrected values is more stable across batches (Figure 2D). Moreover, we found that 1.3% of the DEGs identified using the uncorrected values showed opposite fold changes in the different sample subset comparisons, which we attribute to batch effects. Comparatively, only 0.02% of the DEGs identified using the corrected values show such opposite fold changes. We next examined the difference in DEG overlap for each cell type using the F-score (Figure 2E). We found that for all cell types, the corrected values gave higher DEG overlaps in the subset comparisons. This higher overlap is particularly stronger among certain similar cell types such as the different B cell subtypes. This ability to identify subtle cell differences is especially attractive for big data integration. To check the reliability of the DEGs, we also computed the DEGs between all pairs of cell types using the batch-corrected and raw values of all 50 samples, and compared them with the DEGs computed within individual samples. Assuming that the true DEGs can also be found in most individual samples, we counted the frequency of conserved DEGs being found in the DEGs identified within individual samples. A higher frequency denotes higher reliability for the detected DEGs. We found the DEGs identified with batch-corrected values to be more conserved in each single sample than the DEGs identified with raw values (Figure 2F).

Using batch corrected values for DEG identification is highly controversial with the choice of uncorrected values being commonly stated as the preferred approach. In this study, we demonstrated how batch effects can negatively affect the use of uncorrected values for DEG identification with cell types not found in all batches. By using batch corrected values, we reduced the number of DEGs with incorrectly identified fold change directions and increased the overall accuracy. For cell types found in all batches, the choice of uncorrected versus corrected values should not matter, as the presence of the cells across all batches should cancel out the batch effects. Another commonly encountered scenario in DEG identification is that of technical batch effects confounding with biological or patient effects. While the former is nuisance and should be removed, the latter can be of significance to the study at hand. The disentanglement of both effects remains an unsolved challenge when all batches do not have matching biological samples to serve as a reference to guide the data integration. As such, uncorrected values remain the only choice for investigating biological effects present between samples.

To test the impact of batch effects on TF activity prediction, we applied SCENIC^15^ to the raw and batch-corrected data to predict the TF activities of individual cells. In the TF activity generated UMAP for uncorrected values, the batch effects are clearly present among the same cell types across different batches (Figure 2G). For the FastIntegration’s corrected values, most of the batch effects was eliminated with the same cell types forming their respective clusters and the batches mixed within.

### FastIntegration allows users filter and download public single-cell data in a convenient way

While there are many databases that host single-cell datasets, they possess various drawbacks that make them difficult to exploit for multi-sample integrated analyses. Firstly, sample associated metadata are typically inconsistent across studies. Secondly, different studies use different versions of the reference genome, making it challenging for integration. To solve these problems, we developed the DISCO single-cell database in 2021 to host harmonized single-cell data from different public data sources. Here, we provide functions in FastIntegration that enable users to programmatically access DISCO’s hosted data. With FastIntegration’s API, users can filter samples by various criteria, like tissue, disease, and platform. Moreover, users can also filter the data at cell level by providing expressed or/and unexpressed genes. Combined with the FastIntegration algorithm, users can now explore the DISCO hosted public datasets and integrate them with their own in-house data conveniently and offline.

## Conclusion

Here, we introduced FastIntegration for integrating large single-cell RNA-seq datasets and outputting batch corrected gene expression values. We demonstrated its capacity for large scale batch integration with 4 million cells in 48 hours of runtime through good multicore scaling. With large datasets, it achieved good performance for both homogeneous and heterogeneous data integration when compared with other methods. The batch corrected values produced by FastIntegration can be used for downstream analyses including DEG analysis and regulatory network inference. It also offers programmatic access to DISCO’s hosted single-cell RNA-seq datasets. With the continued growth in single-cell RNA-seq data, FastIntegration will be a valuable tool for large scale data integration tasks.

## Method

### Optimization of Seurat framework for big data integration

Seurat (v3 and v4) integrates data in three steps (anchor identification, sample tree construction, and stepwise batch effect correction). We optimized the pipeline and code of each step to speed them up and reduce memory usage. Although Seurat provide a parallelized version of “FindIntegrationAnchors” using the “future” framework for the anchor identification step, it is memory intensive as it initially loads the data of all batches into memory and R’s garbage collection for parallelization is very slow. To solve this problem, FastIntegration first stores each sample in a separate file and loads them as required. For parallelization, we use the “pbmcapply” package which is comparatively lighter and more user-friendly than “future” framework.

Seurat constructs the sample tree using an iterative function which has an n^2^ x (time to filter anchor table) time complexity where n is the number of samples/batches. The time complexity for filtering the anchor table is also O(n^2^). Therefore, the time complexity for this step is about O(n^4^). To speed up it, FastIntegration converts the anchor table into a “data.table” object which has a key and index for each data row. The time complexity for filtering the “data.table” object is therefore O(log n). To calculate the similarities between samples, FastIntgeration uses the “group_by” functions which have a O(n) time complexity. Thus, the resulting time complexity of our sample tree construction in FastIntegration is reduced to O(nlog(n)).

At the integration stage, Seurat integrates input samples according to the sample tree and needs n-1 iterations (n is the number of samples). At each iteration, PCA is performed on the query data to find the k nearest neighbors (KNN) of each query cell in the PCA space. These operations become the processing bottleneck for big data integration. While the motivation for doing PCA in each round of integration is to get the precise KNN graph, we reasoned that the nearest neighbors are most likely to fall within the same unintegrated dataset as the query cell. Thus, FastIntgeration only runs PCA once on all unintegrated datasets and identifies the nearest anchors in this PCA space, which is used for the subsequent integration. This allows the calculation of integrated expression values for different genes to be parallelized, accelerating integration. Another problem in Seurat’s integration scheme is the risk of over- or mis-correcting gene expression values. To alleviate this problem, in each round of integration, the corrected gene expression value will be set to 0 if it is less than the minimal value of uncorrected value, or set to the maximal value of uncorrected value if it is greater than the maximal value of uncorrected value. Furthermore, Seurat computes a fixed number of k neighbors for each cell to construct the weight matrix of anchors while FastIntegration fits a Gaussian distribution to the distances of k neighbors and removes neighbors with a Z score great than 3. With these modifications, FastIntgeration can remove outlier gene expression values and keep the sparsity of data, avoiding problem of long vector being unsupported in large sparse matrices.

### Assessment of memory usage and runtime

We compared the memory usage and CPU runtime of FastIntegration with Seurat. For a fair comparison, we also enabled parallelization in Seurat using 150 threads. We then tested their performance on different sample sizes ranging from 10 to 200. To obtain the runtime, we used the command “time -v” to evaluate. To obtain the total memory usage by all threads, we used an in-house script to get the maximum memory usage of each job. We first cleared all jobs in the server and then recorded the used memory at every second during the run and determined the peak memory usage after the job finished. All jobs were run on a Linux server with 2x AMD^®^ EPYC 7552 CPUs at 2.20GHz, 2□TB of DDR4 memory, and an 8T SSD disk.

### Benchmark of data integration

We compared FastIntegration with four state-of-the-art methods, BBKNN, Harmony, Scanorama, and scVI. We used two different datasets that represent two common scenarios. The first dataset is a homogeneous dataset which contains 235,886 cells from 50 blood samples, and the second is a heterogeneous dataset with 235,886 cells from 50 blood samples and 32,582 cells from 10 lung samples. We used two evaluation metrics to compare the local and global performance of each method. The local level metric used was the Local inverse Simpson’s index (LISI) by Korsunsky et al (https://github.com/immunogenomics/LISI). We calculated the cell type LISI (cLISI) and batch LISI (bLISI) scores of the integration results from each method and used the F1 score^16^ of cLISI and bLISI as a combined assessment of cell type purity and batch mixing. We also used the Adjusted Rand Index (ARI) to evaluate the global integration performance. We annotated each sample manually and calculated the ARI between cell clusters and cell type annotation. To avoid the influence of clustering resolution, we tested a range of resolutions (0.1 to 1) and used the maximum ARI achieved. The first 50 PCs were used as input to BBKNN and Harmony, and 3000 features selected by FindIntegrationFeatures were used as input to FastIntegration, Scanorama, and scVI. The number of epochs was set to 100 for scVI. All other parameters were set as defaults.

### Differential gene expression analysis and single-cell regulatory network inference

The dataset of 50 blood samples was used to test the performance of DEGs identification and regulatory network inference. We first removed cell types with fewer than 50 cells or greater than 30% in only one sample. After that, 253 cell type pairs were retained for DEGs identification. For each cell type pair, we identified the DEGs twice. The first batch of DEGs was identified between cell type 1 in samples 1-25 and cell type 2 in samples 26-50 and the second batch was identified between cell type 1 in samples 26-50 and cell type 2 in samples 1-25. The “FindMarkers” function in the Seurat package was used for DEG identification and only DEGs with an q value less than 0.05 were retained for further analyses. The F-score (measuring accuracy) is defined as (TP□+□TN)/(TP□+□TN□+□FP□+□FN), which was also used in our previous benchmarking paper^10^. SCENIC was used to infer the single-cell regulatory network and predicted TF activity. The AUC scores produced by SCENIC were used to do cell clustering.

## Supporting information

clusters showing high mutual ARI scores

while the major cell types were well separated, confirmed by their marker gene expressions

with FastIntegration ranked second for both homogenous and heterogeneous integration

FastIntegration were able to maintain the neutrophils as a separate cluster

integrate the immune cells from both blood and lung samples

## Declarations

### Ethics approval and consent to participate

Not applicable.

### Consent for publication

Not applicable.

### Author contribution

Jinmiao Chen conceptualized and supervised the study. Mengwei Li, Xiaomeng Zhang developed the method and analyzed the data. Mengwei Li generated the figures and tables. Kok Siong Ang, Jinmiao Chen, Mengwei Li and Xiaomeng Zhang wrote the manuscript.

### Availability of data and materials

The codes of FastIntegration are maintained in the GitHub: https://github.com/JinmiaoChenLab/FastIntegration

## Acknowledgements

The authors would like to thank the handling editor and anonymous reviewers for their constructive comments.

## Additional files

**Additional file 1: Table S1: Detailed description of datasets.**

**Additional file 2: Figure S1: Comparison FastIntegration and Seurat for integrating 20 blood samples.** A) UMAP of data integrated by Seurat and FastIntegration. B) The mutual ARI between clusters in the data integrated using Seurat and FastIntegration. C) ARI between cell type annotations and clusters at different resolution.

**Additional file 3: Figure S2: UMAP plot of COVID-19 data coloured by cell type and gene expression of selected markers.**

**Additional file 4: Figure S3: 1/cLISI and sLISI of different methods.** Cell type LISI and sample LISI scores computed for A) homogenous data integration and B) heterogeneous data integration.

**Additional file 5: Figure S4: Visualisation of homogeneous data integration by different methods.** UMAP of homogeneous data integration by the different integration methods, colored by A) cell type B) study. Neutrophils are highlighted using red circles.

**Additional file 6: Figure S5: Visualisation of heterogeneous data integration by different methods.** UMAP of heterogeneous data integration by the different integration methods, colored by A) dataset B) cell type.

**Additional file 7: Figure S6: UMAPs of heterogeneous data integration by different integration methods, colored by tissue.**

