## Supplementary figures and images for "FastIntegration: a versatile R package for accessing and integrating large-scale single-cell RNA-seq data"

### clusters showing high mutual ARI scores

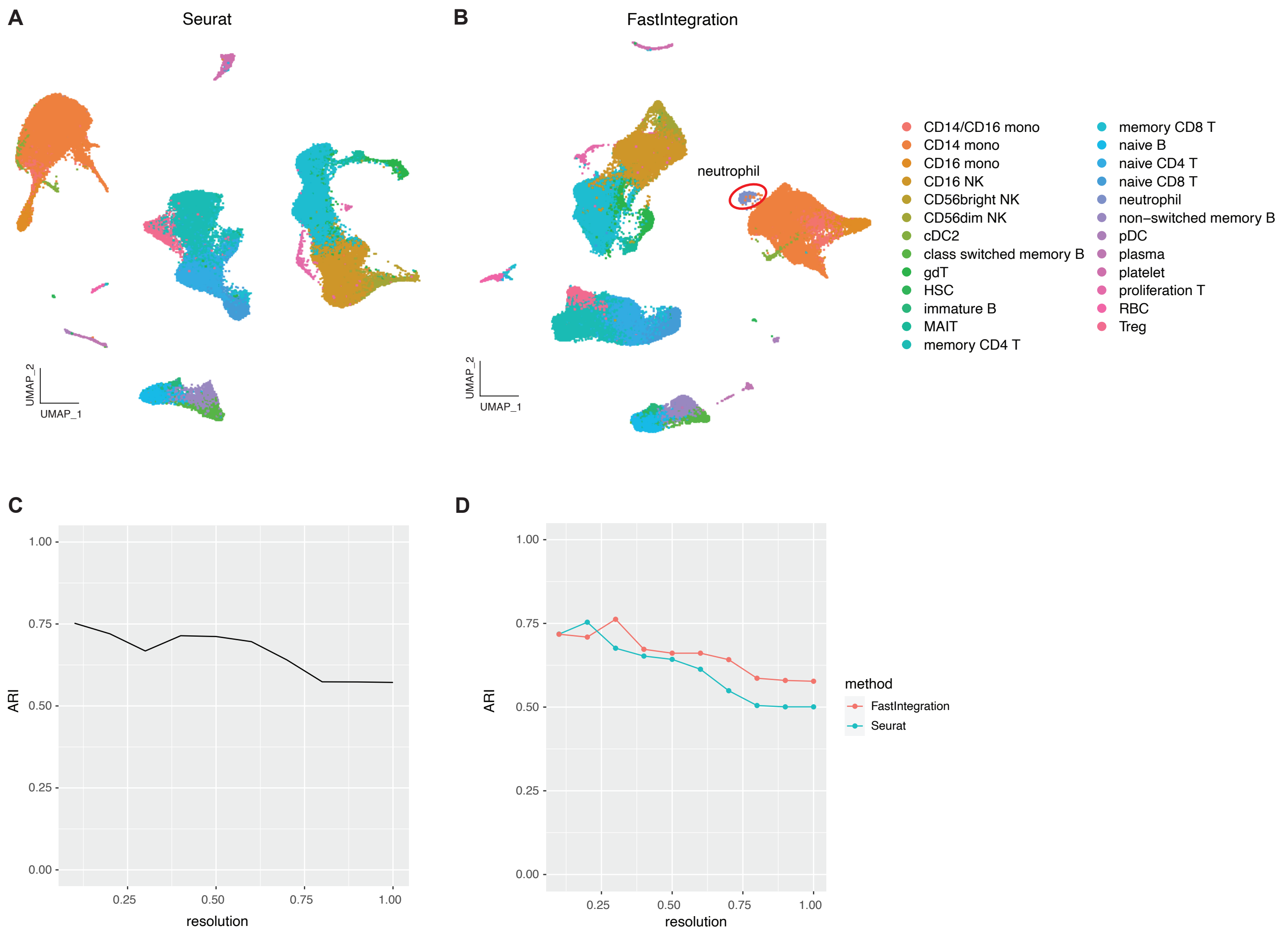

### FastIntegration were able to maintain the neutrophils as a separate cluster

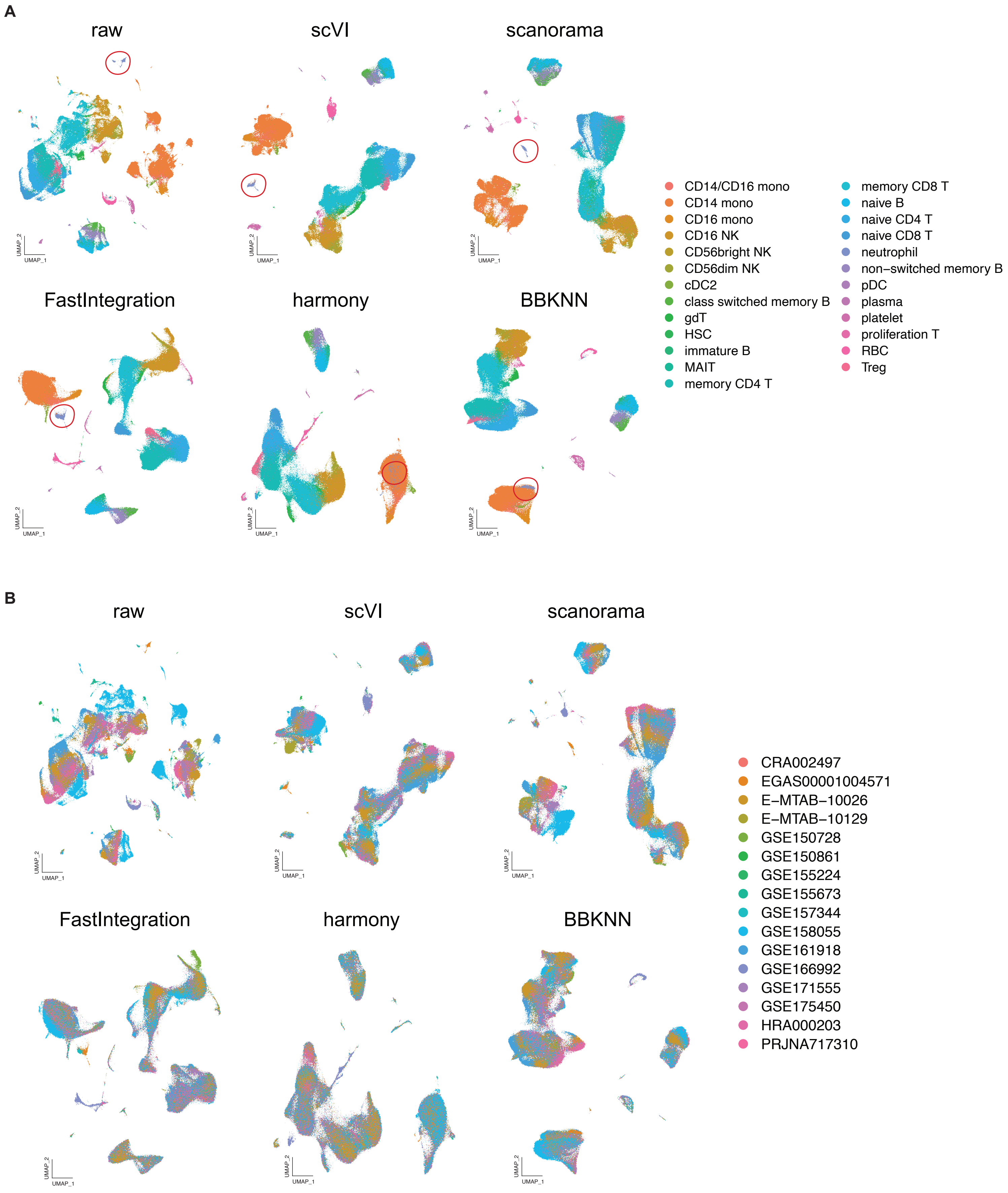

### integrate the immune cells from both blood and lung samples

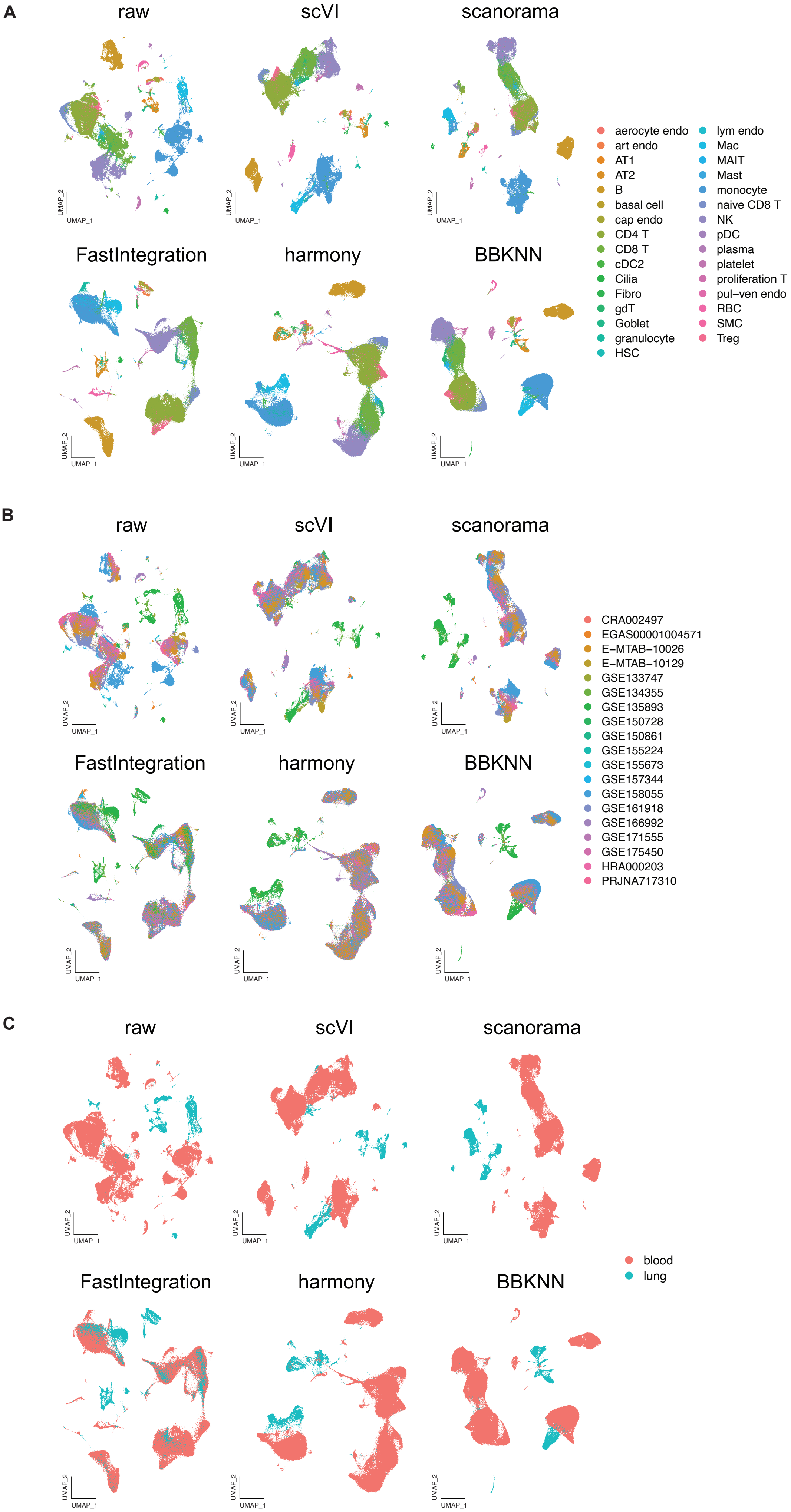

### while the major cell types were well separated, confirmed by their marker gene expressions

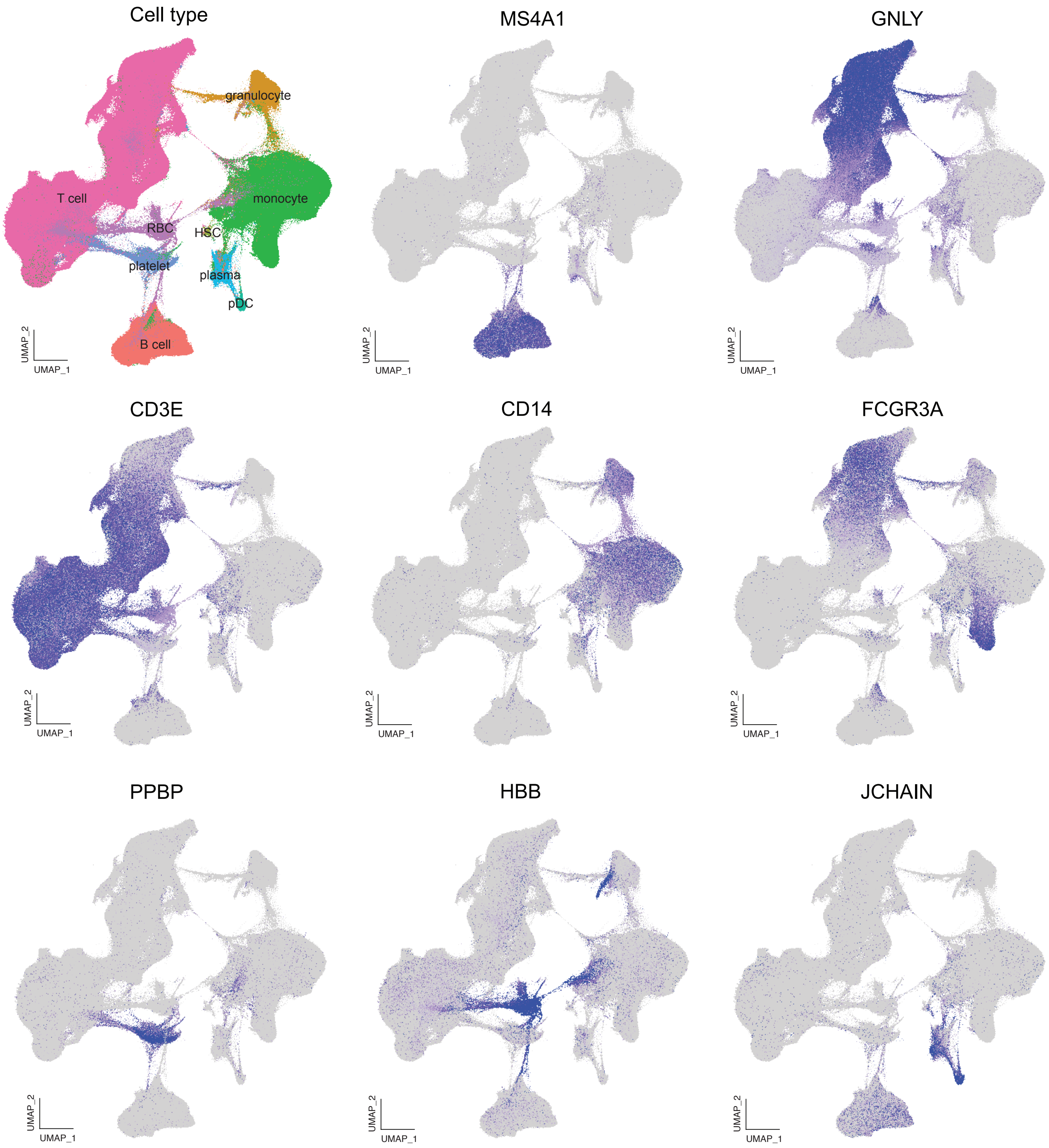

### with FastIntegration ranked second for both homogenous and heterogeneous integration

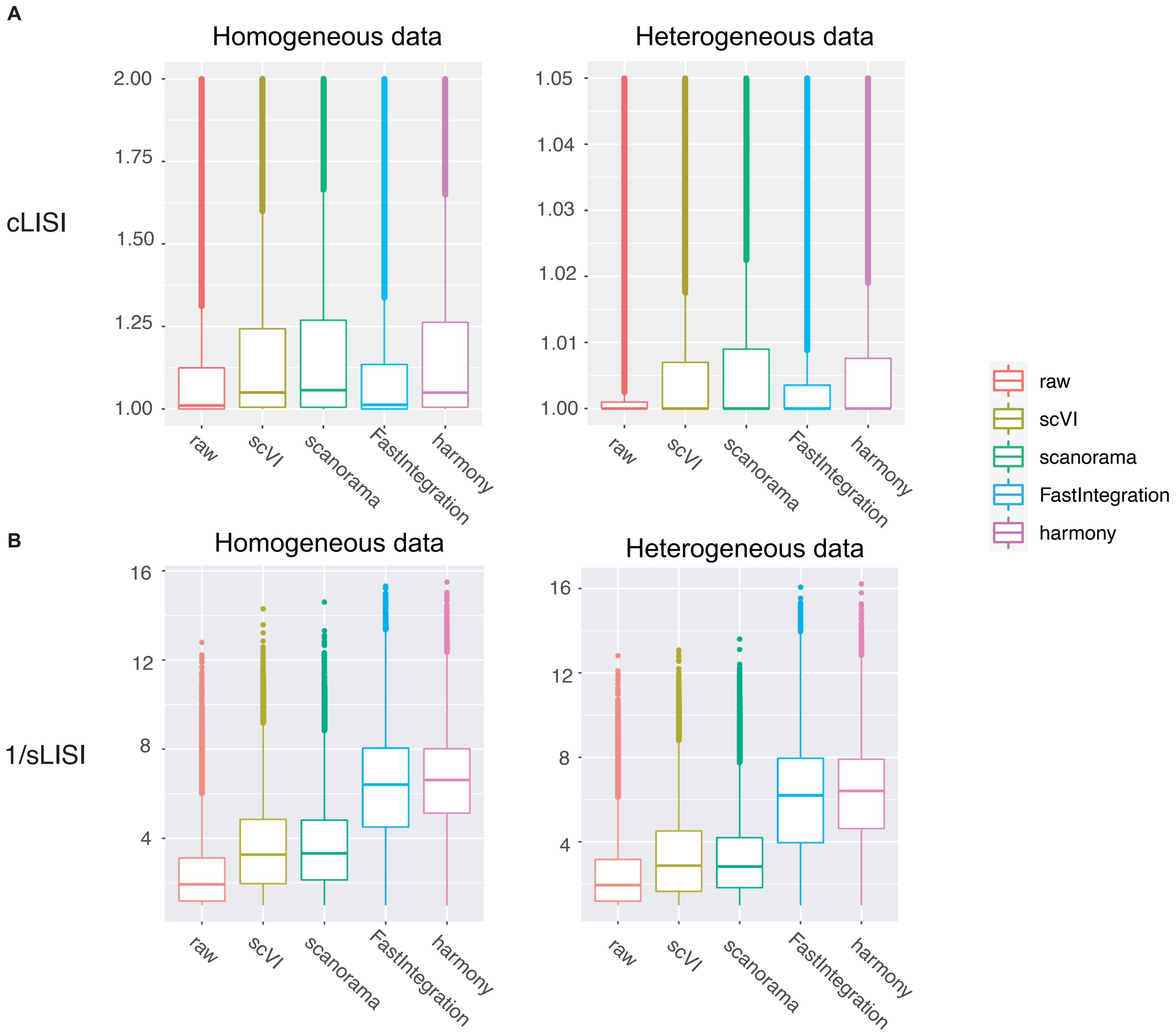
